# Biodiversity loss under future global socio-economic and climate scenarios

**DOI:** 10.1101/235705

**Authors:** Abhishek Chaudhary, Arne O. Mooers

**Affiliations:** Institute of Food, Nutrition and Health, ETH Zurich, Schmelzbergstrasse 9, Zurich 8092, Switzerland; Department of Biological Sciences and IRMACS, Simon Fraser University, Burnaby, British Columbia, Canada V5A1S6

**Keywords:** biodiversity, evolutionary history, future pathways, habitat loss, land use, species extinctions

## Abstract

Efficient forward-looking mitigation measures are needed to halt the global biodiversity decline. These require spatially explicit scenarios of expected changes in multiple indicators of biodiversity under future socio-economic and environmental conditions. Here we link five future (2050 and 2100) global gridded maps (0.25° × 0.25° resolution) available from the land use harmonization (LUH) database that represent alternative representative concentration and shared socio-economic pathways (RCP-SSP) with the countryside species-area relationship model to project the future land use change driven rates of species extinctions and phylogenetic diversity loss (in million years) for mammals, birds and amphibians in each of the 804 terrestrial ecoregions and 176 countries and compare them to the current (1900-2015) and past (850-1900) rates of biodiversity loss. Future land-use changes are projected to commit an additional 209-818 endemic species and 1190-4402 million years of evolutionary history to extinction by 2100 depending upon the scenario, equivalent to 20–80% of the number committed to extinction under current (2015) land use extent. Results show that hotspots of future biodiversity loss differ depending upon the scenario, taxon and metric considered. The most aggressive climate mitigation scenario (RCP2.6 SSP-1), representing a world shifting towards a radically more sustainable path including increasing crop yields, reduced meat production and reduced tropical deforestation coupled with high trade, projects the lowest land use change driven global biodiversity loss followed by RCP8.5 SSP-5, RCP6.0 SSP-4 and RCP7.0 SSP-3. Interestingly, the scenario with the second most aggressive climate target (RCP3.4 SSP-4) projected the highest biodiversity loss among the five scenarios tested. This is because it represents a world with continued high consumption in rich countries and increased land clearing for crop production in species rich, low-income countries such as Indonesia, Madagascar, Tanzania, Philippines and DR Congo. These contrasting results illustrate that the strategies to prevent climate change could simultaneously contribute to reduction in current high rates of biodiversity loss, but only if habitat preservation is incorporated into national and global sustainable development plans.

## Introduction

The rapid decline in global biodiversity has major consequences for ecosystem functioning and human wellbeing [1] and therefore, international agreements such as the United Nations Sustainable Development Goals (SDGs; [2]) and the Convention on Biological Diversity Aichi targets [3] commit to reducing these losses. The Intergovernmental Platform on Biodiversity and Ecosystem Services (IPBES) has identified that reporting past, present and future trends of biodiversity at global and regional levels and development of scenarios is key to help decision makers evaluate different policy options [4,5].

Importantly, habitat destruction and degradation due to human land use for commodity production is, and is expected to remain, the major driver of biodiversity loss [6]. It is therefore surprising that a recent literature review [7] concluded that biodiversity scenarios over the past 25 years have focused on the future direct impacts of climate change, with only a few exploring the biodiversity outcomes due to future land use change. Moreover, most biodiversity scenarios have focused on changes in species richness as the only indicator of biodiversity change [8,9]. To meet the call of the IPBES and to project a more accurate picture of the future of biodiversity, we need advancements in both land use projections and in models that translate these projections onto changes in different aspects of biodiversity (e.g. taxonomic, genetic, functional). We present attempts to do both here.

Land-Use Harmonization products (LUH-1; [10]) delivered with the IPCC Fifth Assessment Report opened up new opportunities for exploring the impacts of a range of possible land-use trajectories on biodiversity. These products connect future scenarios calculated by multiple Integrated Assessment Models (IAMs; [11]) with historical land use data into a single consistent, spatially gridded set of land-use change scenarios. They provide annual fractions of five land uses (primary vegetation, secondary vegetation, pasture, cropland and urban) at the 0.5° × 0.5° scale between 1500 and 2100 for four Representative Concentration Pathway (RCP) scenarios [12]. Each RCP describes an alternative future climate scenario with a specific radiative forcing (global warming) target (e.g. 2.6, 3.4, 4.5, 6.0 or 8.5 W/m^2^) to be reached by the end of the century through the adoption of mitigation efforts [9,11]. Radiative forcing under each scenario is often considered as a proxy for the expected amount of atmospheric warming [13].

However, the main limitations of above LUH-1 land-use datasets is their relatively coarse spatial resolution (0.5 degree) and a limited number of land use categories (five) per grid cell. Such simplified representation of land use is due primarily to the computational complexity of the models underlying these datasets [7]. For accurately predicting impacts on biodiversity, both the location and intensity of future land-use change projections are important because certain regions host a disproportionately high number of endemic species [14] and because the responses of species can differ widely depending upon management intensity [15].

Recently, a new set of future scenarios has been developed using the so called scenario matrix architecture approach [16]. These scenarios are based on the combination of RCPs (describing carbon emissions trajectories, [12]), and shared socioeconomic pathways (SSPs), that describe development and socio-economic trajectories [17,18]. Five SSPs have been developed (SSP-1 to SSP-5) providing different trajectories of future socio-economic development, from most to least benign, and include possible trends in population, income, agriculture production, food & feed demand, global trade, as well as land use [19]. Within each SSP, climate policies such as afforestation can be introduced in order to reach a particular radiative forcing level target consistent with a RCP [20].

In 2017, as a part of the Coupled Model Intercomparison Project (CMIP6; [21]), the updated land use harmonization dataset (LUH2 v2f) for the period 2015-2100 was released (http://luh.umd.edu/data.shtml) providing annual gridded fractions of 12 land-use types at 0.25° x 0.25° resolution under five scenarios varying in climate target (RCP) and Shared Socioeconomic Pathways (SSPs). The dataset also provides annual land use maps for the past (850-2014). This is a major improvement over the previous coarse scale and simplified LUH1 dataset [11] and provides an excellent opportunity to evaluate the past extinction rates as well as the future biodiversity trajectories under different RCP-SSP scenario combinations. With regard to biodiversity models translating the land use change into biodiversity loss, previous studies [8,9] have employed the classic form of species-area relationship model (SAR) that assumes that no species can survive in any human land use, thereby potentially overestimating species loss. Chaudhary & Brooks [22] showed that the countryside SAR model [23,24], which accounts for the fact that some species are tolerant to human land uses better predicts habitat loss driven extinctions than classic SAR on a global scale. Chaudhary and Brooks [22] proposed and tested a novel approach to parameterize the countryside SAR by leveraging the species-specific habitat classification scheme database [25] and projected the species extinctions (mammals, birds, amphibians) due to land use change to date in 804 terrestrial ecoregions [26]. Another advantage of the countryside SAR model is that it allows allocating the total extinctions to individual land uses in the regions and thereby enables the identification of major drivers of species loss [22,27,28].

It has been argued that in addition to species richness, phylogenetic diversity (PD; also referred to as evolutionary history) may be a useful indicator of the biodiversity value of a region (see, e.g. [29-31]) because PD represents the evolutionary information within the set of species, and a higher PD may offer a region with more functional diversity via complementarity, resilience, and more options to respond to global changes [30, 32-34].

Unlike species richness [8,9], no study to date has evaluated the loss of phylogenetic diversity under future global scenarios. This is primarily because the complete dated species-level phylogenies for large taxonomic groups [31,35,36] and straightforward methods that can translate species extinctions in a region into the loss of PD from the evolutionary tree in which these species are found, have only recently become available [37,38]. This is relevant because previous studies have shown that regions with high projected species extinctions might not overlap with regions projected with high PD loss [37].

Here we project potential future extinctions, and phylogenetic diversity (PD) loss, for mammals, birds and amphibians in each of the 804 terrestrial ecoregions (and 176 encompassing countries) associated with land-use changes to 2050 and 2100 under five alternative scenarios, and compare them to projected extinctions under current (2015) and past (850, 1900) land use extent. For comparison purposes, we also calculate the rate at which species are getting committed to extinction in the past (850-1900), present (1900-2015) and future time slices (2015-2050 & 2015-2100) by dividing additional projected extinctions with the time interval (e.g. 85 years for 2015-2100) and the total species richness of the taxon (see methods Eq. 2, 5).

We feed the future gridded maps generated by the five RCP-SSP combination scenarios (RCP 2.6 SSP-1, RCP 7.0 SSP-3, RCP 3.4 SSP-4, RCP 6.0 SSP-4, and RCP 8.5 SSP-5) available from recent LUH2 dataset into the newly parameterized countryside SAR model (Chaudhary & Brooks [22]) to project number of endemic species committed to extinction in each ecoregion. To project the associated evolutionary history of the three taxa committed to extinction in each ecoregion and country (in million years), we apply the novel approach recently proposed by Chaudhary et al. [37] that combines countryside SAR, species-specific evolutionary distinctiveness (ED) scores [39,31] and a linear relationship between the cumulative ED loss and PD loss derived through pruning simulations on global evolutionary trees [37]. We identify hotspot ecoregions and countries projected to suffer high biodiversity loss under each future scenario and also allocate the total extinctions and PD loss to individual human land use types to identify the major drivers in each ecoregion and country.

We found that the most aggressive climate change mitigation scenario coupled with a sustainable socio-economic trajectory (RCP2.6 SSP-1) projects the least global biodiversity loss in future, demonstrating that global climate and biodiversity goals can be aligned. However, the second-most aggressive climate change mitigation scenario in a world characterized by increasing disparities across countries (RCP3.4 SSP-4) projects the highest biodiversity loss, due to accelerated clearing of natural vegetation in species rich, low-income tropical countries [19]. These results highlight that strategies to mitigate climate change (e.g. replacing fossils with fuel from bioenergy crops) need to be accompanied by sustainable socio-economic pathways such as strong land use change regulations, reduced human consumption, high global trade and technological innovations in order to ensure a win-win situation for global climate, biodiversity and sustainable development goals.

## Results

### Future natural habitat loss

Across all five coupled RCP-SSP scenarios, an additional 5.87–11.77×10^6^ km^2^ of natural vegetative cover loss is projected between 2015 and 2050 to make way for human land uses, equivalent to 10–20% of total remaining area of natural vegetation across all ecoregions in 2015. From 2050-2100, a further 5.4-12.1 ×10^6^ km^2^ natural vegetation loss is projected, depending upon the scenario. Among the five scenarios, RCP 2.6 SSP-1 projected the least loss of natural vegetative cover followed by RCP 8.5 SSP-5, RCP 6.0 SSP-4 and RCP 7.0. SSP-3. The scenario with second most stringent climate mitigation target, which is coupled with a less benign shared social pathway (RCP 3.4 SSP-4) projected the greatest loss of natural vegetative cover.

Secondary vegetation (e.g. logged forests) will be responsible for the conversion of the majority of natural vegetation area from 2015-2100, followed by conversion to cropland. While currently in 2015, the combined area of pasture and rangelands for livestock grazing constitutes the most dominant human land use, all five scenarios project that by 2100, secondary vegetation will be most dominant human land use type globally. The RCP2.6 SSP-1 and RCP8.5 SSP-5 both project abandonment of grazing area from 2015-2100 (with subsequent increases in secondary vegetation) while the other three scenarios project a marginal increase.

### Projected species richness and evolutionary history loss

In Table 1, we infer that a total of 1023 endemic species (199 mammals, 222 birds and 602 amphibians) are committed to extinction (or have gone extinct) due to land use change to date (2015). The 95% confidence interval taking into account the uncertainty in the slope of the species-area relationship (z-values; [22]) is 958-1113 projected endemic extinctions globally (Supplementary Table S3).

**Table 1.** Projected number of endemic species (Eq. 1) and phylogenetic diversity (Eq. 4; PD in million years) committed to extinction under past (850, 1900 AD), present (2015) and future (2050, 2100) land use extent. M, B, A and T correspond to mammals, birds, amphibians and total respectively. See methods for details on five climate mitigation (RCP) and socio-economic (SSP) scenario combinations. Supplementary Table S3 presents the 95% confidence interval for the estimates.

| Year | Scenario | Projected endemic extinctions |  |  |  | Projected PD loss (MY) |  |  |  |
| --- | --- | --- | --- | --- | --- | --- | --- | --- | --- |
|  |  | M | B | A | T | M | B | A | T |
| 850 AD | Past | 6 | 5 | 15 | 25 | 0 | 3 | 20 | 24 |
| 1900 AD | Past | 74 | 86 | 212 | 372 | 268 | 271 | 1137 | 1675 |
| 2015 AD | Present | 199 | 222 | 602 | 1023 | 948 | 828 | 3493 | 5270 |
| 2050 AD | RCP2.6 SSP-1 | 219 | 236 | 665 | 1120 | 1039 | 872 | 3875 | 5787 |
|  | RCP7.0 SSP-3 | 249 | 260 | 746 | 1255 | 1202 | 976 | 4394 | 6572 |
|  | RCP3.4 SSP-4 | 267 | 270 | 764 | 1301 | 1338 | 1022 | 4517 | 6876 |
|  | RCP6.0 SSP-4 | 252 | 253 | 735 | 1241 | 1272 | 943 | 4391 | 6606 |
|  | RCP8.5 SSP-5 | 241 | 255 | 747 | 1244 | 1157 | 920 | 4421 | 6499 |
| 2100 AD | RCP2.6 SSP-1 | 241 | 256 | 734 | 1232 | 1226 | 941 | 4293 | 6459 |
|  | RCP7.0 SSP-3 | 302 | 301 | 883 | 1485 | 1523 | 1150 | 5203 | 7876 |
|  | RCP3.4 SSP-4 | 398 | 408 | 1035 | 1841 | 1997 | 1575 | 6100 | 9672 |
|  | RCP6.0 SSP-4 | 320 | 319 | 883 | 1522 | 1581 | 1203 | 5273 | 8057 |
|  | RCP8.5 SSP-5 | 278 | 281 | 825 | 1385 | 1365 | 1043 | 4878 | 7286 |

This number is equivalent to ~30% of all endemic species across all ecoregions and corresponds to a projected loss of 5270 million years (MY) of phylogenetic diversity (95% confidence interval: 3912-6967 MY). The taxonomic breakdown is 948, 828 and 3493 MY of phylogenetic diversity (PD) loss for mammals, birds and amphibians respectively. Compared to inferences for 850 and 1900, the current total represents a threefold increase in number of species and evolutionary history committed to extinction globally (Supplementary Table S3).

Our projected species extinctions due to current land use extent (2015) for each taxon per ecoregion compare well with the number species documented as threatened with extinction (i.e. those with threat status of critically endangered, endangered, vulnerable, extinct in wild or extinct) by the IUCN Red List in 2015 [14], with correlation coefficients of 0.74, 0.75, and 0.85 for mammals, birds and amphibians respectively (see Table S2 and Fig S1 in supporting information for goodness of fit values and additional notes).

As a consequence of further natural land clearing and land use changes from 2015-2100, we project that an additional 209-818 endemic species will be committed to extinction by 2100 (Table 1). The concomitant PD committed to extinction amounts to 1190-4402 million years. Compared to mammals and birds, a three times higher loss of amphibian species is projected owing to their small range and high level of endemism. The highest biodiversity loss is projected to occur under RCP3.4 SSP-4 scenario, followed by RCP6.0 SSP-4, RCP7.0 SSP-3, and RCP8.5 SSP-5 (Table 1). The RCP2.6 SSP-1 scenario is projected to result in least number of additional species and PD committed to extinction by 2100.

### Land use drivers of biodiversity loss

Table 2 shows the contribution of 10 human land use types to the total biodiversity loss (mammals, birds and amphibians combined) in the past, present and future, calculated through allocation factor (0≤ *a_g,i,j_*≤1) in Eq. 6, 7 (methods). Currently in 2015, the global secondary vegetation (forests + non-forests), grazing land (pasture + rangeland) and cropland are responsible for 378, 333 and 289 projected extinctions representing 37%, 33% and 28% of the total 1023 projected extinctions respectively.

**Table 2.** Contribution of different human land use types (in %; Eq. 6, 7) to total number of species and phylogenetic diversity (mammals, birds and amphibians combined) committed to extinction in the past (850, 1900 AD), present (2015) and future (2100) under five climate mitigation (RCP) and socio-economic (SSP) scenario combinations.*

| Land use type | Past |  | Present<br>2015AD | % of total projected extinctions in 2100AD |  |  |  |  |
| --- | --- | --- | --- | --- | --- | --- | --- | --- |
|  | 850 | 1900 |  | RCP2.6 | RCP7.0 | RCP3.4 | RCP6.0 | RCP8.5 |
|  | AD | AD |  | SSP-1 | SSP-3 | SSP-4 | SSP-4 | SSP-5 |
| Secondary Veg. (forests) | 0 | 28 | 28 | 35 | 28 | 21 | 30 | 35 |
| Secondary Veg. (non-forests) | 0 | 19 | 9 | 13 | 8 | 8 | 8 | 11 |
| Pasture | 7 | 13 | 21 | 11 | 17 | 15 | 20 | 16 |
| Rangeland | 44 | 16 | 11 | 8 | 10 | 8 | 10 | 8 |
| Urban | 0 | 0 | 2 | 3 | 3 | 3 | 4 | 3 |
| C3 annual crops | 20 | 9 | 11 | 9 | 13 | 11 | 11 | 10 |
| C3 permanent crops | 13 | 7 | 9 | 11 | 10 | 13 | 8 | 8 |
| C4 annual crops | 9 | 4 | 5 | 3 | 6 | 4 | 4 | 6 |
| C4 permanent crops | 2 | 1 | 1 | 5 | 1 | 13 | 3 | 1 |
| C3 Nitrogen fixing crops | 4 | 2 | 2 | 2 | 3 | 2 | 2 | 3 |

\*Note that the contribution % remains the same for both metrics (species extinctions and evolutionary history) since the same allocation factor is used (Eq. 6).

By 2100, the contribution of grazing area to total extinctions is projected to decrease to 19 - 30% depending upon the scenario, while secondary vegetation will be responsible for the majority of projected extinctions under four out of five scenarios (Table 2). The exception is SSP-4 RCP3.4 scenario, where agriculture land (particularly C3 and C4 permanent crop area) is projected to be the most damaging land use type contributing to 48% of all projected extinctions in 2100.

### Rates of biodiversity loss

We found that for all three taxa combined, the current projected rate (over the period 1900-2015) of endemic species committed to extinction is 242 (95% confidence interval: 228-261) extinctions per million species years (E/MSY; Eq. 2) which is ~20 times higher than the inferred past rate (850-1900) of projected extinctions (Fig 1a). The rate of current phylogenetic diversity (PD) loss is 156 (95% confidence interval: 124-185) million years per million phylogenetic years (MY/MPY; Eq. 5) that is also ~20 times higher than the past rate of 8 MY/MPY (Fig 1b). Amphibians have highest rates of loss (515 E/MSY, 289 MY/MPY) followed by mammals (192, 120) and birds (106, 60). Supplementary Table S3 presents the taxon-specific (mammals, birds and amphibians) rates of projected extinctions for past, present and future.

**Fig 1.**
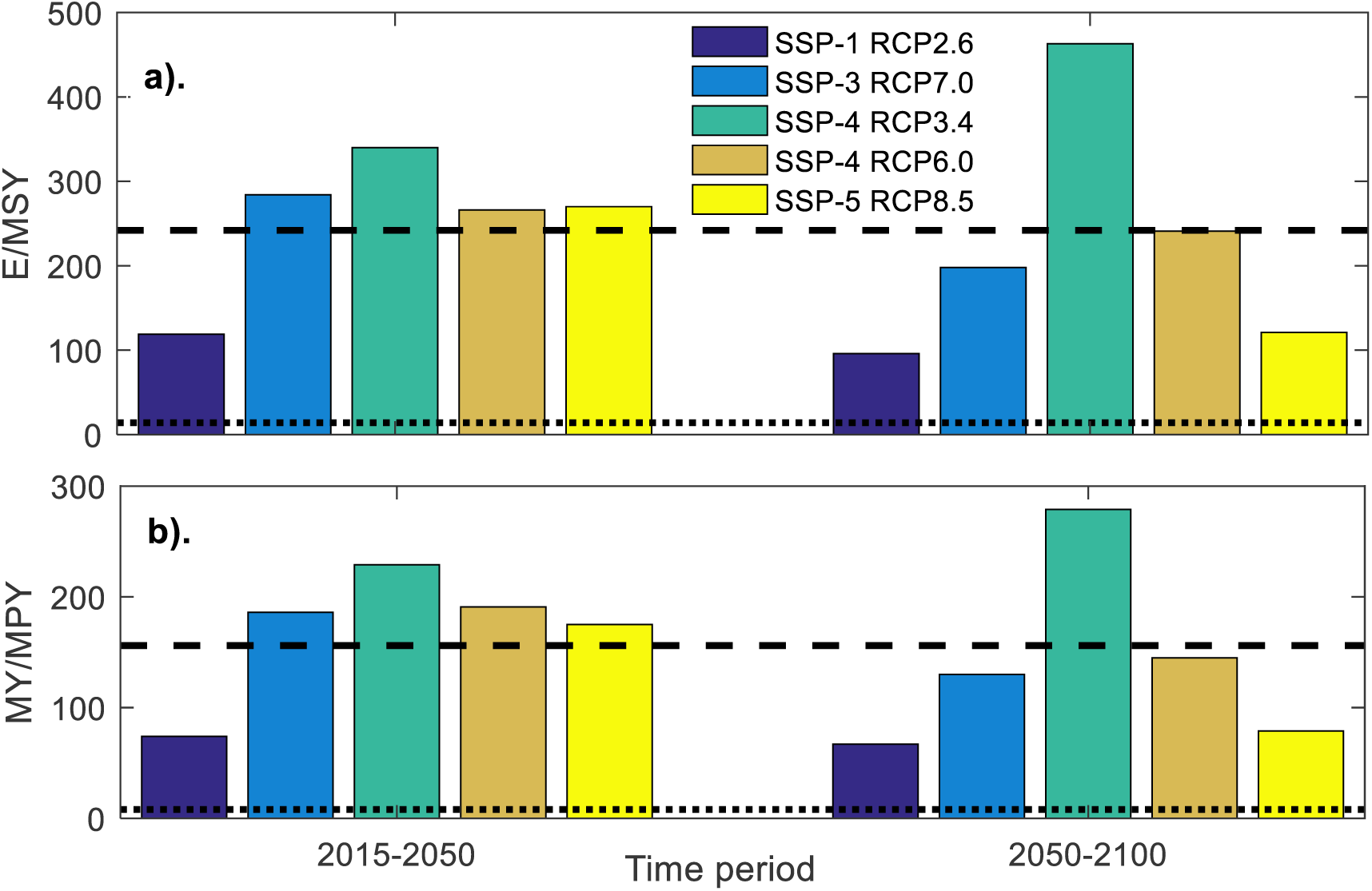
Past, present and future rates of projected biodiversity loss. Future rate of biodiversity loss (mammals, birds, amphibians combined) in projected endemic species extinctions per million species years (a) and projected PD loss per million phylogenetic years (b) under five climate mitigation (RCP) and socio-economic (SSP) scenario combinations. For comparison, past (850-1900) rate of loss is represented by the dotted line and current (1900-2015) rate of loss is shown by a dashed line. See Supplementary Table S3 for taxon-specific (amphibians, mammals, birds) rates of loss along with the 95% confidence intervals for the estimates.

The rate of biodiversity loss is projected to increase for the period 2015-2050 under all scenarios but one: under the combination of the lowest climate forcing (RCP2.6) and the most benign developmental path (SSP-1), biodiversity loss is expected to reduce to half its current level. This projected amelioration extends to other scenarios over the period 2050-2100: the rate of biodiversity loss will reduce to the levels below the current rates under all scenarios except RCP3.4 SSP-4 (under which it will increase to twice its current levels and ~1.5 times the rate in 2015-2050 period; see Fig 1). Compared to 2015-2050 period, we found a huge reduction in rate of biodiversity loss in the second half of the century under the RCP8.5 SSP-5 and RCP7.0 SSP-3 scenarios (a reduction of 149 E/MSY and 86 E/MSY respectively; Fig 1).

### Hotspots of biodiversity loss

Figure 2a shows the current hotspot ecoregions where the land use change to date is projected to cause high amounts of PD loss (Eq. 4). These constitute ecoregions spanning Madagascar, Eastern arc forests (Kenya, Tanzania), northwest Andes (Central Colombia), Appalachian Blue Ridge forests (eastern USA), Peruvian Yungas, Bahia coastal forests (Eastern Brazil) and Palawan forests (Philippines).

**Fig 2.**
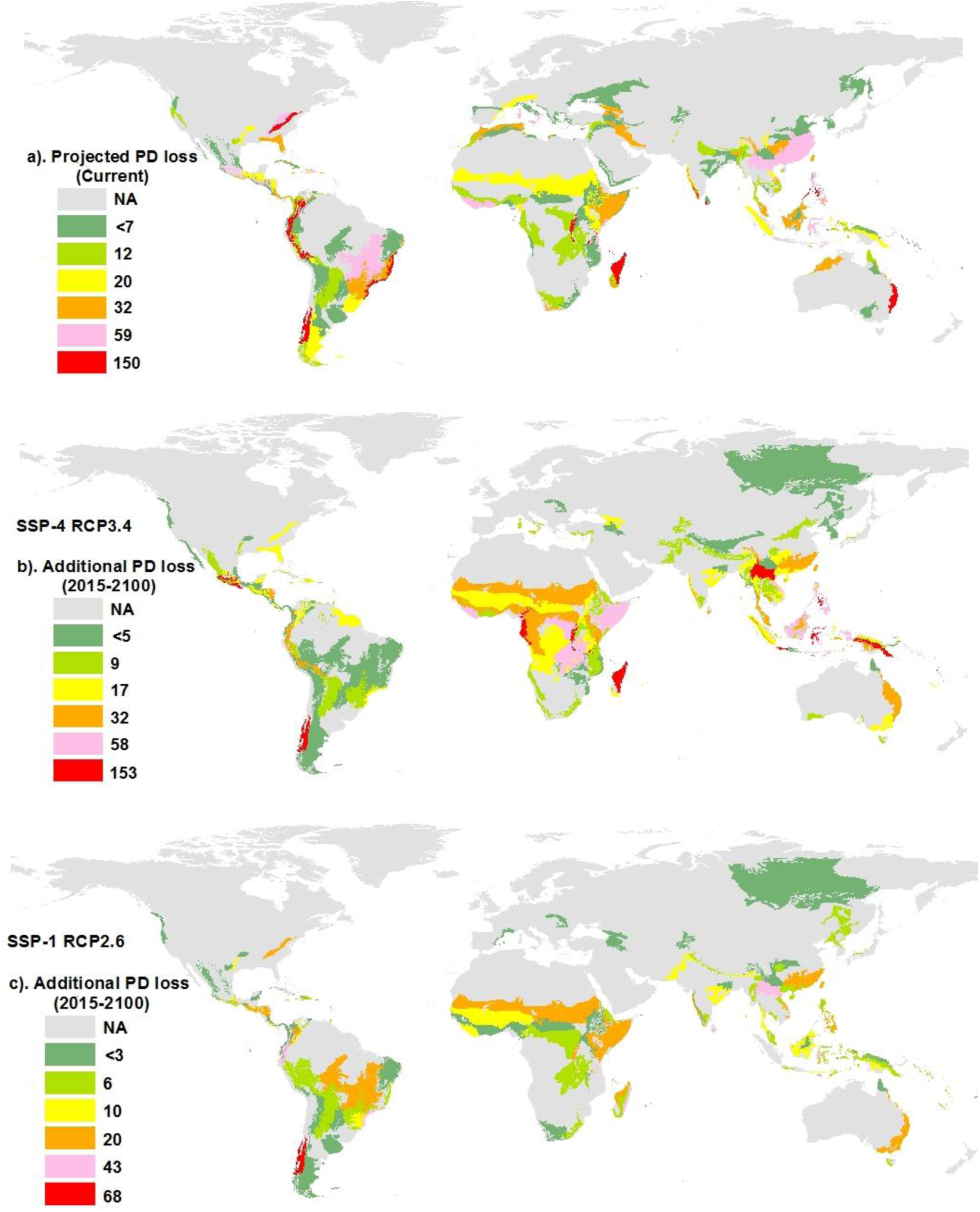
Hotspots of phylogenetic diversity (PD) loss under current and future human land use extent. a). Projected loss of PD in millions of years (MY) associated with projected endemic species extinctions (mammals, birds and amphibians combined) due to current (2015) human land use in each of 804 terrestrial ecoregions, b). Additional PD (in MY) committed to extinction due to land use change under RCP3.4 SSP-4 scenario (worst case) in the period 2015-2100, c). Additional PD (in MY) committed to extinction due to land use change under RCP2.6 SSP-1 scenario (best case) in the period 2015-2100.

In terms of our metrics of biodiversity loss, we found that hotspots of projected species loss do not always overlap with hotspots of projected PD loss. For example, while currently (year 2015) the Appalachian forest ecoregion in eastern USA ranks 11^th^ among the 804 ecoregions in terms of projected species loss (15 extinctions), it ranks 4^th^ in terms of projected PD loss (113 MY), due to its harboring of many old endemic amphibian lineages. Conversely, the Comoros forest ecoregion between Madagascar and East Africa ranks 4^th^ in terms of projected species loss (22 extinctions) but 19^th^ for PD loss – here, endemic species are relatively young.

Among the five future scenarios, RCP3.4 SSP-4 represents the worst case outcome for biodiversity, with an additional 818 species committed to extinction globally (representing 4402 MY of PD) compared to 2015 levels, primarily due to expansion of C3 and C4 permanent crop area (Table 2). Figure 2b shows the additional PD loss per ecoregion due to future land use change (2015-2100) under the RCP3.4 SSP-4 scenario. Most current hotspots of species and PD loss such as Madagascar and Eastern arc forests are projected to continue losing the biodiversity under this scenario. However, other current hotspots of loss, such as the Appalachian forests (USA) and the Bahia forest (Brazil) ecoregions, are not expected to experience significant land use change, and hence no additional biodiversity loss is projected under this scenario.

Importantly, due to clearing of natural vegetation under the RCP3.4 SSP-4 scenario, new hotspots of global species and PD loss emerge, i.e. ecoregions that have hitherto been relatively undamaged from human land use. This includes Sulawesi montane forests (Indonesia), Cameroonian highland forests, and central range montane forests (Papua New Guinea, Indonesia), each with an additional 20-30 projected species loss and around 100 MY of additional PD loss (Fig 2b). Supplementary Table S4 presents the additional species and PD loss in each of 804 ecoregions under all future scenarios.

The RCP2.6 SSP-1 scenario represents the best case in terms of future outcomes for biodiversity. Compared to the worst case scenario above (RCP3.4 SSP-4), 75% fewer species extinctions and evolutionary history loss are projected under scenario for the period 2015-2100 (Table 1). Under the other three scenarios (RCP7.0 SSP-3, RCP6.0 SSP-4 and RCP8.5 SSP-5), the projected biodiversity loss in the period 2015-2100 is about half of that under the worst case and twice of that under the best case (Table 1).

We found that hotspots of future biodiversity loss differ depending upon the scenario. For example, while the RCP3.4 SSP-4 scenario projects an additional 11 MY of PD loss in Serra do Mar forest ecoregion in eastern Brazil over the period 2015-2100, the projected loss is three times higher (34 MY) under the RCP2.6 SSP-1 scenario. Conversely, unlike RCP3.4 SSP-4, RCP2.6 SSP-1 scenario projects almost no additional biodiversity loss in Cameroonian highland or central range forest ecoregions.

Hotspots of future biodiversity loss also differ depending upon the taxon. For example, under the worst case for biodiversity scenario (RCP3.4 SSP-4), the ecoregion with highest number of projected additional endemic mammal species extinctions (over 2015-2100) is the Sulawesi rain forest ecoregion in Indonesia. The Galapagos Island ecoregion is projected to bear the most additional bird species extinctions while the Madagascar lowland forests will see the most amphibian species extinctions.

We also calculated the projected biodiversity loss for each of the 176 countries as well as the contribution of each land use type to total projected loss in each nation under all future scenarios. Supplementary Table S5 presents the additional biodiversity loss in each country and supplementary Table S6 presents the biodiversity loss allocated to each of the 10 land use types in each country under all future scenarios. Table 3 shows the additional biodiversity loss projected for seven World Bank regions under five future scenarios (derived by summing up the projected species and PD loss estimate for all countries within a region). As expected, the majority of future biodiversity loss is projected to occur due to land use change in tropical countries with a high endemism per unit area. The projected biodiversity loss is small in North America, Europe & Central Asia (EU&CA), Middle-East and North Africa (MENA) regardless of scenario (Table 3), while the amount of biodiversity loss in the other four regions differ depending upon the scenarios. For example in the period 2015-2050, the RCP7.0 SSP-3 scenario projects the highest endemic extinctions in Sub-Saharan Africa (SSA), while the RCP3.6 SSP-4 projects the highest losses in East Asia & Pacific (EAP) and the RCP8.5 SSP-5 scenario highlights Latin America & Caribbean (LAC; Table 3).

**Table 3.** Additional numbers of endemic species (SR) and PD (in millions of years) committed to extinction (for mammals, birds and amphibians combined) in seven World Bank regions for the periods 2015-2050 and 2050-2100 due to land use change under five coupled climate mitigation (RCP) and socioeconomic (SSP) scenarios. Bold figures represent the regions with highest projected loss for a particular scenario. See supplementary Table S5 for country-specific numbers under each scenario.

| Metric | Period | Scenario | EAP | EU&CA | LAC | MENA | N.America | S.Asia | SSA |
| --- | --- | --- | --- | --- | --- | --- | --- | --- | --- |
| SR | 2015-2050 | SSP-1 RCP2.6 | **30** | 0 | 28 | 0 | 3 | 14 | 13 |
|  |  | SSP-3 RCP7.0 | 58 | 0 | 59 | 0 | 3 | 27 | **77** |
|  |  | SSP-4 RCP3.4 | **96** | 0 | 49 | 0 | 4 | 32 | 88 |
|  |  | SSP-4 RCP6.0 | 55 | 0 | 40 | 0 | 4 | 27 | **82** |
|  |  | SSP-5 RCP8.5 | 50 | 0 | **66** | 0 | 3 | 29 | 62 |
|  | 2050-2100 | SSP-1 RCP2.6 | **45** | 1 | 41 | 0 | 2 | 8 | 17 |
|  |  | SSP-3 RCP7.0 | 65 | 2 | 63 | 0 | 1 | 3 | **95** |
|  |  | SSP-4 RCP3.4 | **281** | 7 | 100 | 1 | 1 | 7 | 137 |
|  |  | SSP-4 RCP6.0 | 101 | 4 | 42 | 2 | 0 | 6 | **127** |
|  |  | SSP-5 RCP8.5 | **65** | 3 | 31 | 0 | 1 | 4 | 43 |
| PD | 2015-2050 | SSP-1 RCP2.6 | 139 | 5 | **167** | 0 | 16 | 117 | 73 |
|  |  | SSP-3 RCP7.0 | 257 | 16 | 338 | 1 | 16 | 184 | **482** |
|  |  | SSP-4 RCP3.4 | 535 | 10 | 281 | 1 | 23 | 206 | **544** |
|  |  | SSP-4 RCP6.0 | 330 | 5 | 268 | 1 | 24 | 186 | **518** |
|  |  | SSP-5 RCP8.5 | 286 | 5 | **363** | 1 | 15 | 203 | 354 |
|  | 2050-2100 | SSP-1 RCP2.6 | **269** | 0 | 232 | 1 | 10 | 27 | 130 |
|  |  | SSP-3 RCP7.0 | 403 | 8 | 335 | 1 | 2 | 6 | **557** |
|  |  | SSP-4 RCP3.4 | **1358** | 44 | 513 | 1 | 12 | 50 | 784 |
|  |  | SSP-4 RCP6.0 | 494 | 28 | 179 | 1 | 2 | 26 | **695** |
|  |  | SSP-5 RCP8.5 | **306** | 23 | 169 | 0 | 1 | 8 | 275 |

\*The seven World Bank regions are: East Asia and Pacific (EAP), Europe and Central Asia (EU&CA), Latin America and Caribbean (LAC), Middle-east and North Africa (MENA), North America, South-Asia and Sub-Saharan Africa (SSA).

These varying results also reflect the temporal trajectories of land use change in a particular region under a specific scenario. For example, the RCP8.5 SSP-5 scenario projects almost similar biodiversity loss in LAC (Latin America & Caribbean) and SSA (Sub-Saharan Africa) for the period 2015-2050 but projects substantially higher loss in EAP (East Asia & Pacific) in the period 2050-2100 than the other two regions.

In terms of country level numbers, while RCP3.6 SSP-4 projects just five additional endemic species committed to extinction in Brazil between 2015-2100, the corresponding additional extinctions under RCP8.5 SSP-5 is very high at 42 (supplementary Table S5). Interestingly, additional biodiversity loss in Papua New Guinea over the period 2015-2100 under RCP3.6 SSP-4 is >5 times higher than projected under RCP6.0 SSP-4 scenario. This shows that even under the similar socio-economic trajectory (SSP-4), the impacts may vary within a country due to land use changes that are driven by climate mitigation efforts (RCP3.4 vs. RCP6.0).

Finally, hotspots of future biodiversity loss also differ depending upon the metric of biodiversity loss. For example, between 2015-2050 the most optimistic RCP2.6 SSP-1 scenario would commit the East Asia & Pacific (EAP) region to the greatest number of endemic species extinction, whereas Latin America & Caribbean (LAC) emerges as the region projected to lose the most additional PD (see bold numbers in Table 3).

## Discussion

The large variations in the magnitude and location of projected land use changes and the consequent biodiversity outcomes across different SSP-RCP scenarios and across taxa (Table 1-3; Fig. 1, 2; Supplementary Tables S3-S6) are a result of the complex interplay of mitigation measures adopted to achieve the climate target under a particular RCP [12], the integrated assessment model used for simulation [11], the future socio-economic conditions under each SSP (e.g. land use change regulations, food demand, dietary patterns, global trade, technological change in agriculture sector etc. [17,19]) and the biodiversity theatre on which all this plays out. The ambitious RCP2.6 SSP-1 scenario is projected to result in lowest land use change driven global biodiversity loss. The RCP 2.6 target would be achieved by the deployment of bioenergy (C4 permanent) crops in conjunction with carbon capture and storage technology [40]. While this does entail clearing of some natural habitat and therefore some biodiversity loss, the demand for agricultural products is lowest among the SSPs, due to lower projected population growth, adoption of sustainable food consumption practices and an increase in agricultural yields and global trade [19,41]. These SSP factors outweigh the natural habitat clearing needed to achieve RCP2.6 climate mitigation target, with the net result being relatively low biodiversity loss.

In contrast, the SSP-4 RCP 3.4 (the worst case scenario for projected biodiversity loss) has the climate mitigation measures (deployment of bioenergy crops) and SSP factors (high population growth, lower crop yields and weak land use change regulation in the tropical countries, [42]) working synergistically and leading to large amounts of natural habitat loss in biodiversity hotspots, and consequent biodiversity loss. The RCP6.0 SSP-4 performs better than the RCP3.4 counterpart because the less ambitious climate target necessitates relatively low levels of bioenergy crop expansion.

The RCP8.5 SSP-5 scenario showed the second lowest biodiversity loss despite being characterized by high food waste and diets high in animal source food. This is because of an absence of any explicit climate mitigation efforts that result in land use change, and in addition strong increase in crop yields, global trade and medium levels of land use change regulation [43]. In particular, the land demand and rate of biodiversity loss is substantially reduced in the second half of the century (Fig 1) as the population decreases, consumption levels stabilize and livestock production shifts from extensive to more intensive animal husbandry systems [43]. We note here that our extinction projections do not include losses due to the direct effects of climate change - it may be that the increased global forcing of RCP8.5 counteracts any biodiversity savings due to SSP-5 driven habitat sparing (see below).

Among the five scenarios, RCP7.0 SSP-3 ranks in the middle for biodiversity losses. As the climate mitigation target is not very stringent, the land use change due to bioenergy crop expansion is low. However, socio-economic (SSP-3) factors such as continued high demand for agricultural and animal based products, low agricultural intensification and low trade levels, and no regulations on tropical deforestation, lead to high increase in pasture, and cropland areas for food and feed production [44]. This cascades into high losses of natural vegetation and endemic species extinctions.

Application of countryside SAR allowed us to allocate the total projected loss to individual land use types (Eq. 6, 7; Table 2; Table S6). All future scenarios show an increase in secondary vegetation area at the cost of natural habitat primarily to meet the increasing wood demand [9]. We found that this leads to substantial biodiversity loss in all five scenarios (Table 2) indicating that, regardless of climate mitigation, sustainable forest management will be critical for future biodiversity conservation. This lends support to the call for low-intensity wood harvesting techniques such as reduced impact logging, to protect biodiversity (see, e.g., [15,45]).

Our projections of biodiversity loss are conservative for several reasons. First, the SAR approach we use cannot quantify instances where the land use change wipes out the habitat of *non*-endemic species from all the ecoregions that they occur in [22]. Second, we only considered the species loss for three taxa (mammals, birds and amphibians) for which necessary data were available for this analysis, and we do not know how to scale patterns from these three taxa to biodiversity in general [46]. Third, and importantly, we did not include direct climate-change driven extinctions. Climate change can lead to loss of habitat for several species and drive them to extinction [47-50]. That said, direct climate-change driven biodiversity loss is also expected to be least under the best scenario we used here, RCP2.6 SSP-1. Including additional drivers such as habitat fragmentation [51], overexploitation/hunting, invasive species and pollution would likely increase all biodiversity loss estimates [6], though perhaps differentially under the various SSPs.

Overall, our results corroborate previous findings that land use change over coming decades is expected to cause substantial biodiversity loss. For example, Van Vuuren et al. [8] used four Millennium Ecosystem Assessment scenarios [52] and projected that habitat loss by year 2050 will result in a loss of global vascular plant diversity by 7–24% relative to 1995 using the classic form of species area-relationship model [53]. Jetz et al. [54] found that at least 900 bird species are projected to suffer >50% range reductions by the year 2100 due to climate and land cover change under four Millennium Ecosystem Assessment scenarios. Jantz et al. [9] applied classic species-area relationship model to gridded land-use change data associated with four RCP scenarios (without the SSPs, [10]) and found that by 2100 an additional 26-58% of natural habitat will be converted to human land uses across 34 biodiversity hotspots and as a consequence, about 220 to 21000 additional species (plants, mammals, birds, amphibians and reptiles) will be committed to extinction relative to 2005 levels. More recently, Newbold et al. [55] used four RCP scenarios (again without the SSPs) and generalized linear mixed effects modeling to project that local plot-scale species richness will fall by a further 3.4% from current levels globally by 2100 under a business-as-usual land-use scenario; with losses concentrated in low-income, biodiverse countries.

## Conclusions

This study is the first, to our knowledge, to employ a newly parameterized countryside species-area relationship model and species-specific evolutionary isolation scores with specific land use projections to calculate the past, present and future rates of biodiversity loss through two indicators (species richness and phylogenetic diversity) across all ecoregions and countries. Results show that the current rate (1900-2015) of projected biodiversity loss is ~20 times the past rate (8501900) and is set to first increase in the period 2015-2050 under all scenarios (except under RCP2.6 SSP-1) and then decrease to levels below the current rate in the period 2050-2100 (expect under RCP3.4 SSP-4).

We found that out of the five future scenarios, the one most aggressive in terms of climate change mitigation effort (RCP2.6 SSP-1) is also the one projected to result in lowest land use change driven global biodiversity loss because of adoption of sustainable path to global socio-economic development. However, the poor performance of the RCP3.4 SSP-4 scenario demonstrates the need for caution: strategies to mitigate climate change (e.g. replacing fossils with fuel from bioenergy crops) are likely to result in adverse global biodiversity outcomes if they involve clearing of natural habitat in the tropics.

Even in the best case RCP2.6 SSP-1 scenario, more than 10 million km^2^ of primary habitat is projected to be converted into secondary vegetation for wood production or into permanent crops for bioenergy production in developing countries by the year 2100, potentially committing an additional 200 species and 1000 MY of evolutionary history to extinction. This implies that if we accept the importance of biodiversity to human well-being [56], then even a dramatic shift towards sustainable pathways such as healthy diets, low waste, reduced meat consumption, increasing crop yields, reduced tropical deforestation and high trade, e.g. as specified under the RCP2.6 SSP-1 scenario is not likely not enough to fully safeguard its future. Additional measures should focus on keeping the natural habitat intact through regulating land use change in species rich areas [57], reducing the impact at currently managed areas through adoption of biodiversity-friendly forestry/agriculture practices [15,45] or restoration efforts [58], and controlling the underlying drivers such as human consumption to reduce land demand [59].

We identified hotspots of biodiversity loss under current and alternative future scenarios and note that these hotspots of future biodiversity loss differ depending upon the scenario, taxon and metric considered (Table 3; Fig. 2; Table S4, S5). This lends support to calls to carry out multi-indicator analyses in order to get a more comprehensive picture of biodiversity change [60].

Overall, the quantitative information we present here should inform the production and implementation of conservation actions. Combining multiple threats (i.e. climate, habitat loss, direct human pressure and fragmentation) with the coupled RCP-SSP scenarios should allow more accurate predictions of biodiversity change. Given that we can allocate these predictions to particular land uses at the country level, incorporating country-level funding and development information (e.g. Waldron et al. [61]) is a logical next step. Such country-level projections should allow better allocation of conservation resources, necessary to move us towards UN *Aichi* Target 13 (preserving genetic diversity) and UN SDG 15 (conserving terrestrial biodiversity).

## Materials and methods

### Future scenarios

Future land use maps at an annual interval (2015-2100) derived using five combinations of shared socio-economic pathways (SSPs) and representative concentration pathways (RCP) that are currently available at LUH2 website (http://luh.umd.edu/data.shtml): RCP 2.6 SSP-1, RCP 7.0 SSP-3, RCP 3.4 SSP-4, RCP 6.0 SSP-4, and RCP 8.5 SSP-5. The RCP8.5 scenario represents business as usual (highest) global warming whereas the lower RCPs signify mitigation scenarios with lesser predicted warming [12,13,62]. The SSP-1 RCP 2.6 scenario was simulated using the Integrated Model to Assess the Global Environment (IMAGE) [41]. The RCP 3.4 SSP-4 and RCP 6.0 SSP-4 scenarios were developed using the Global Change Assessment Model (GCAM; [63]). The RCP 7.0 SSP-3 and RCP 8.5 SSP-5 scenarios were simulated using the Asia-Pacific Integrated Model (AIM; [44]) and the Regionalized Model of Investments and Development-The Model of Agricultural Production and its Impact on the Environment (REMIND-MAgPIE; [43]) respectively. Land use change under a sixth scenario (RCP 4.5 SSP-2; [64]) is not yet available but will be uploaded at LUH2 website in near future. The SSPs are described as follows:

The SSP-1 (*sustainability* – *taking the green road*) scenario represents a world shifting towards a more sustainable path, characterized by healthy diets, low waste, reduced meat consumption, increasing crop yields, reduced tropical deforestation, and high trade, which, together collectively “respects the environmental boundaries” [41].

The SSP-3 (*regional rivalry* – *a rocky road*) scenario represents a world with resurgent nationalism, increased focus on domestic issues, almost no land use change regulations, stagnant crop yields due to limited technology transfer to developing countries and prevalence of unhealthy diets with high shares of animal based products and high food waste [44].

In SSP-4 (*inequality* – *a road divided*), the disparities increase both across and within countries such that high-income nations have strong land use change regulations and high crop yields while the low-income nations remain relatively unproductive with continued clearing of natural vegetation. Rich elites have high consumption levels while the others have low [42].

The SSP-5 (*fossil fueled development* – *taking the highway*) is characterized by rapid technological progress, increasing crop yields, global trade and competitive markets where unhealthy diets and high food waste prevail. There are medium levels of land use change regulations in place, meaning that tropical deforestation continues, although its rate declines over time [43].

### Land use data

We downloaded from the LUH2 website gridded maps that provide the fraction of each of 12 land use types for each (0.25° × 0.25° resolution) cell for five separate points in time: past (850 and 1900), present (2015), and two future years (2050 and 2100). The 12 land use classes include two classes of primary undisturbed natural vegetation (forests, non-forests) and 10 human land uses (see Table 1). The 10 human land uses comprise two secondary (regenerating) vegetation land use classes (forest, non-forest), two grazing (pasture, rangeland), one urban and five cropland (C3 annual, C3 permanent, C4 annual, C4 permanent and C3 nitrogen fixing crops) uses. C3 and C4 correspond to temperate (cool) season crops and tropical (warm) season crops respectively.

Next, we calculated the area of each land use type in each grid cell by multiplying the fractions with the total cell area. Finally, for each point in time (850, 1900, 2015, 2050, and 2100) and scenario, we calculated the area (km^2^) of each of the 12 land classes in each of the 804 terrestrial ecoregions [26] by overlaying the area maps with ecoregion boundaries in ArcGIS.

### Projecting species extinctions

For each year and scenario, we fed the estimated areas of each of 12 land use types into the countryside species-area relationship (SAR; [23]) to estimate the number of endemic species projected to go extinct (*S_lost_*) due to total human land use within a terrestrial ecoregion *j* as [22]:

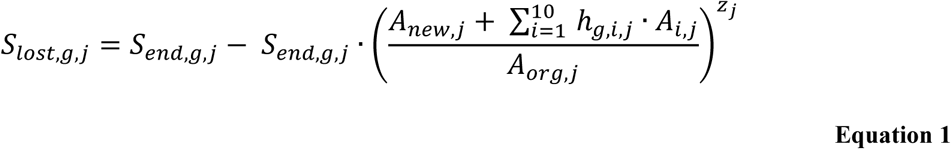

Here *S_end,g,j_* is the number of endemic species of taxon *g* (mammals, birds, amphibians) in ecoregion *j*, *A_org,j_* is the total ecoregion area, *A_new,j_* is the remaining natural habitat area in the ecoregion (primary vegetation forests + primary vegetation non-forests), *h_g,i,j_* is the affinity of taxon *g* to the land use type *i* in the ecoregion *j* (which is based on the endemic species richness of the taxon in that land use type relative to their richness in natural habitat), *A_i,j_* is the area of a particular human land use type *i* (total of 10), and *Z_j_* (z-value) is the SAR exponent. The exponent *z* is the slope of the log-log plot of the power-law SAR describing how rapidly species are lost as habitat is lost [65]. Note that the classic SAR is a special case of the countryside SAR, when *h* = 0.

Following Chaudhary & Brooks [22], we obtained the numbers of endemic species of each taxon per ecoregion (*S_end,g,j_*) from the IUCN species range maps [14], z-values (*z_j_*) from Drakare et al. [65] and the taxon affinities (*h_g,i,j_*) to different land use types from the Habitat Classification Scheme database of the IUCN [25]. This database provides information on the human land use types to which a particular species is tolerant to and within which it has been observed to occur. For each of the 804 ecoregions, we counted the number of endemic species per taxon that are tolerant to the human land use type *i* and divided this by the total number of endemic species of that taxon occurring in the ecoregion, to obtain the fractional species richness. The affinity of taxonomic group *g* to the land use type *i* is then calculated as the proportion of all species that can survive in it (fractional richness), raised to the power 1/*z_j_* [22,23]. We derived the 95% confidence intervals for projected species extinctions in each ecoregion by considering uncertainty in the z-values (*z_j_*) [65]. See supplementary Table S1 for further details and sources of all model parameters.

In order to validate the predicted number of species currently (in 2015) committed to extinction per ecoregion by the parameterized model, we compare them with the documented (“observed”) number of ecoregionally endemic species currently listed as threatened with extinction (i.e. those with threat status of critically endangered, endangered, vulnerable, extinct in wild or extinct) on the 2015 IUCN Red List [14]. Supplementary Table S2 and Chaudhary & Brooks [22] present further details on validation procedure.

We used Eq. (1) to calculate the number of species committed to extinction (*S_lost,g,j_*) in each ecoregion due to human land use extent in five different years: 850, 1900, 2015, 2050 and 2100 (T1 to T5). We then derived the number of additional projected extinctions for the four periods: (850–1900), (1900–2015), (2015–2050) and (2015–2100) by subtracting *S_lost,g,j_* at T1 from *S_lost,g,j_* at T2 and so on.

In addition to absolute projected species loss numbers, we also calculated the rate of projected species loss per taxon for these four time periods by dividing additional extinctions with the time interval and the total species richness of the taxon (*S_org,g_*).

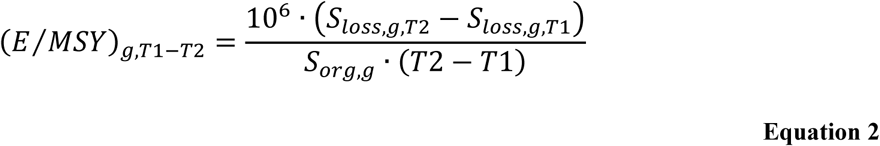

The rate of projected species loss unit is projected extinctions per million species years (E/MSY), representing the fraction of species going extinct over time. We note that this definition of rate of species loss is different than the traditional definition (see e.g. [66,67]) in that it includes both - species going extinct as well as additional species committed to extinction (extinction debt) over the time interval. The traditional definition only considers the extinctions that have been materialized in the time period and does not include the extinction debt.

### Projecting evolutionary history loss

We projected the loss of phylogenetic diversity in each ecoregion in three steps. First, we obtained the evolutionary distinctiveness (ED) scores for all species of birds, mammals, and amphibians from Chaudhary et al. [37]. Next, for each ecoregion *j*, we used a MATLAB routine to sum the ED scores for *m* randomly chosen endemic species from that ecoregion:

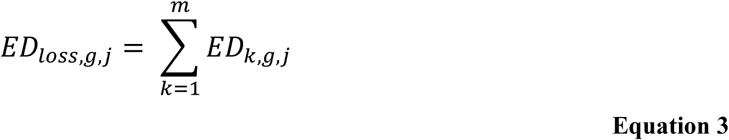

Here *m* is the projected species loss as calculated by the countryside SAR for ecoregion *j* and associated taxa *g* in Eq. 1 (i.e. *m* = *S_lost,g,j_*). We repeated this 1,000 times through Monte Carlo simulation and used the mean and 95% confidence interval of the cumulative ED lost (in million years) per ecoregion.

Finally, we use the least-squares slope (*s_g_*) of the linear model (derived by Chaudhary et al. [37]) describing the tight relationships (R^2^ > 0.98) between ED loss vs. PD loss for each of the three taxa to convert cumulative ED loss per ecoregion to PD loss per ecoregion for a given taxon (Eq. 4). The linear slope (*s_g_*) values used for mammals, birds and amphibians are 0.51, 0. 61 and 0.46 respectively [37]. See Chaudhary et al. [37] for more details on the approach.

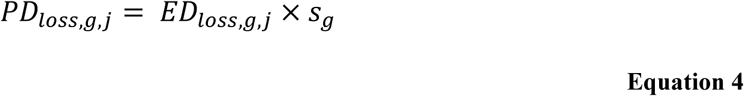

Similar to the rate of species loss (Eq. 2), we also calculated the rate of PD loss in the units million years lost per million phylogenetic years (MY/MPY):

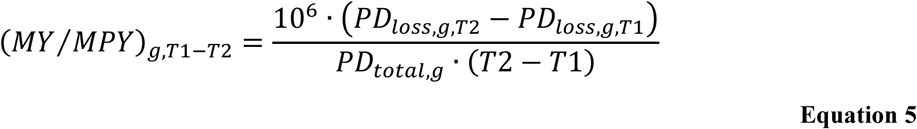

### Drivers of evolutionary history loss in each ecoregion

The projected species extinctions and PD lost per ecoregion is then allocated to the different land use types *i* according to their area share *A_i,j_* in the ecoregion *j* and the affinity (*h_g,i,j_*) of taxa to them. The allocation factor *a_g,i,j_* for each of the 10 human land use types *i* (such that for each ecoregion *j*, 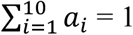) is [22,27,28]:

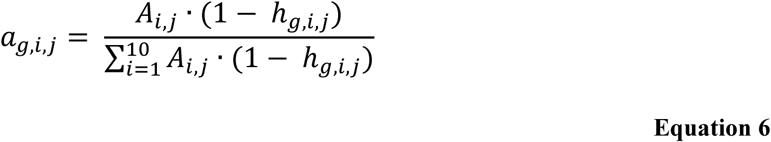

The contribution of different land use types towards the total PD loss in each ecoregion is then given by [37]:

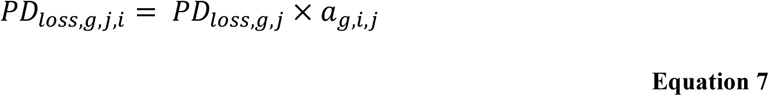

Eq. 7 thus provides the projected PD loss (MY) caused by a particular land use in a particular ecoregion. Replacing *PD_loss,g,j_* with *S_loss,g,j_* provides the contribution of each land use type to total species extinctions per ecoregion [22]. We also calculate the projected extinctions and *PD_loss_* for each of 176 countries based on the share of ecoregion and different land use types within them [22,27,28].

## Acknowledgements

We thank Anthony Waldron, Dan Greenberg, Caroline Tucker and the members of the iDiv working group “Conservation and Phylogenetics” sponsored by the Synthesis Centre of the German Centre for Integrative Biodiversity Research (iDiv) Halle-Jena-Leipzig, for ongoing discussion. AOM acknowledges ongoing funding through Discovery and Accelerator Grants from the Natural Sciences and Engineering Research Council (NSERC) of Canada.

